# Creatine transporter deficient rat model show motor dysfunction, cerebellar alterations and muscle creatine deficiency without muscle atrophy

**DOI:** 10.1101/2021.10.12.464080

**Authors:** Lara Duran-Trio, Gabriella Fernandes-Pires, Jocelyn Grosse, Ines Soro-Arnaiz, Clothilde Roux-Petronelli, Pierre-Alain Binz, Katrien De Bock, Cristina Cudalbu, Carmen Sandi, Olivier Braissant

## Abstract

Creatine (Cr) is a nitrogenous organic acid and plays roles as fast phosphate energy buffer to replenish ATP, osmolyte, antioxidant, neuromodulator, and as a compound with anabolic and ergogenic properties in muscle. Cr is taken from the diet or endogenously synthetized by the enzymes AGAT and GAMT, and specifically taken up by the transporter SLC6A8. Loss-of-function mutations in the genes encoding for the enzymes or the transporter cause Cerebral Creatine Deficiency Syndromes (CCDS). CCDS are characterized by brain Cr deficiency, intellectual disability with severe speech delay, behavioral troubles, epilepsy and motor dysfunction. Among CCDS, the X-linked Cr transporter deficiency (CTD) is the most prevalent with no efficient treatment so far. Different animal models of CTD show reduced brain Cr levels, cognitive deficiencies and together they cover other traits similar to those of patients. However, motor function was poorly explored in CTD models and some controversies in the phenotype exist in comparison with CTD patients. Our recently described *Slc6a8*^*Y389C*^ knock-in (KI) rat model of CTD showed mild impaired motor function, morphological alterations in cerebellum, reduced muscular mass, Cr deficiency and increased guanidinoacetate content in muscle, although no consistent signs of muscle atrophy. Our results indicate that such motor dysfunction co-occurred with both nervous and muscle dysfunction, suggesting that muscle strength and performance as well as neuronal connectivity might be affected by this Cr deficiency in muscle and brain.

## INTRODUCTION

Creatine (Cr) is an organic compound used as fast phosphate energy buffer to replenish ATP by the Cr/phosphocreatine (PCr)/creatine kinase (CK) system, important in tissues with high energy demand such as muscle or brain. Cr is also an osmolyte, an antioxidant, and a suggested neuromodulator^1,2^. Cr is taken from diet or endogenously synthetized in a two-step pathway by the enzymes AGAT (arginine-glycine-amidinotransferase) and GAMT (guanidinoacetate-methyltransferase), and transported into cells by SLC6A8 (also named CRT/CreaT/CT1, henceforth CrT)^3^. Guanidinoacetate (GAA) is the intermediate product of Cr synthesis while creatinine (Crn) is the spontaneous breakdown product of Cr.

Deficits in synthesis or transport of Cr cause Cerebral Creatine Deficiency Syndromes (CCDS)^4-7^. CCDS are characterized by brain Cr deficiency, intellectual disability with severe speech delay, behavioral abnormalities, seizures and motor dysfunction. Among CCDS, CTD is the most frequent, with an estimated prevalence of 2% of males with intellectual disability and, unlike the other two CCDS, with no efficient treatment so far^8^. CTD is an X-linked gene disorder caused by loss-of-function mutations in *SLC6A8*.

Several CTD animal models were generated with ubiquitous deletions^9-11^ or ubiquitous point mutation^12^ in *Slc6a8*. Other conditional, ubiquitous tamoxifen-induced^13^ or brain-specific^14-18^ knock-out (KO) CrT mice were also developed to better understand the role of CrT and provide additional cues about CTD pathology. Rodent CTD models with ubiquitous *Slc6a8* modification exhibit characteristics similar to those of CTD patients (Cr-deficient brain, decreased body weight, cognitive deficits). Together, they cover other features of CTD such as increased urinary Cr/Crn ratio and behavioral abnormalities including autistic-like traits or seizures^9-12,14,16,19,20^. However, despite the known Cr effects in neuron morpho-functional development^21-23^ and its ergogenic and anabolic properties in muscle^24^, motor function was poorly studied in CTD. Only one CTD mouse model described a motor dysfunction phenotype and it was linked with Cr-deficient and metabolically-impaired muscle^11^, although no muscle Cr deficiency was found in the unique CTD patient analyzed for this^25^.

Here we aim to shed light on this controversy and explore motor function in the recently described *Slc6a8*^*Y389C*^ rat model of CTD^12^, taking advantage of the higher proximity of rats over mice to human physiology, biochemistry and behavior^26^.

## METHODS

### Animals

Adult wild-type (WT) and knock-in (KI) *Slc6a8*^*xY389C/y*^ males (3-4 months-age) were used for all experiments; animals were genotyped and housed as previously described^12^. Rats were fed with a Cr-devoid vegetarian diet (Safe 150). All experiments were performed with approval of Canton de Vaud veterinary authorities (auth.VD-3284) in accordance with the Swiss Academy of Medical Science, and followed the ARRIVE Guidelines 2.0. Efforts were made to minimize stress and number of animals used.

### Tissue collection

Animals were anesthetized with 4-5% isoflurane, plasma and urine were collected as described^12^, and animals were sacrificed by rapid decapitation. Brain and quadriceps were rapidly dissected out and prepared for biochemical or histological analysis. Tissues for biochemical analysis were frozen and stored at -80°C. Tissues for histological analysis were fixed in 4% paraformaldehyde (PFA) in PBS overnight at 4°C, then rinsed with PBS and sunk in 18% (muscle) or 30% (brain) sucrose before being embedded in Tissue-Tek (OCT4583, for muscle) or gelatin/sucrose (7.5% gelatin, 15% sucrose in phosphate buffer, for brain), frozen and stored at -80°C.

### Measure of Cr, GAA, CK, Crn and acetylcholine

Plasmatic and urinary Crn was measured as described^12^. Plasmatic CK was measured on a COBAS8000 automate (Roche, Switzerland). For intramuscular measure of Cr and GAA, 3 replicates/animal were used. Powder of frozen muscle was homogenized in H_2_O (sonication, 4°C, 10s) and centrifuged (10’000g, 5min, 4°C). Supernatant was used to measure Cr and GAA by LC-MS/MS as described^12,27^. Muscle acetylcholine (ACh) determination was performed according to instructions (Sigma-Aldrich MAK056), using 3 replicates/animal. Total choline is the sum of free choline and ACh.

### Quantitative PCR

Muscle tissue (20mg) was processed and analyzed as described^60^, with these particularities: 500µl TRIzol, 200µl chloroform, primers listed in **Table 1**, and 2 housekeeping genes (*GAPDH* and *rRNA 18S*) to compensate for variations in mRNA input and efficiency of reverse-transcription.

**Table1.**
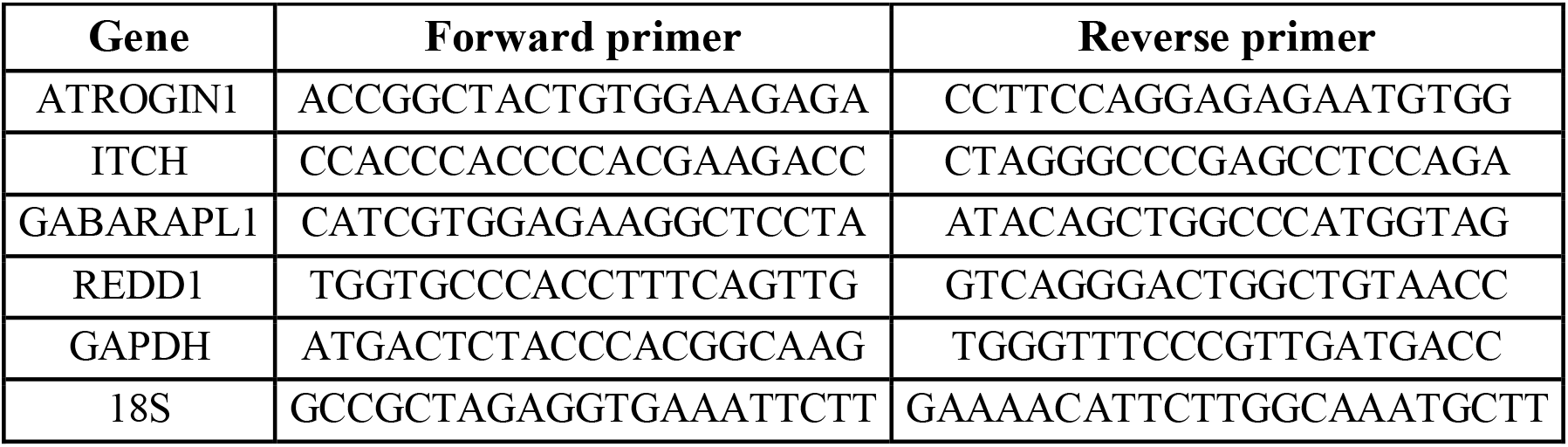
Primers used for qPCR.

### Western Blot

Muscle tissue (20 mg) was processed and analyzed by western blotting (10µg of total protein) as described^60^. Primary and secondary antibodies (listed in supplementary materials) were used for chemiluminescent detection of proteins. Membranes were scanned with chemidoc imaging system (Bio-rad) and quantified (Image lab software, Bio-rad).

### Histology

Cryopreserved tissue was cut with a cryostat (Leica; 20µm thickness). Slices were mounted on microscope slides, warmed (∼40°C, 20min), tempered and frozen at -80°C until use for immunofluorescence or hematoxylin-eosin staining.

#### Immunofluorescence

Slides were tempered 15min at room temperature (RT), incubated with 4%PFA-PBS (30min, RT), washed (PBS, 3×5min) and proceeded for immunofluorescence as decribed^61^. Primary (1:100, overnight 4°C) and secondary (1:200, 1h RT) antibodies are listed in supplementary materials. At the end of immunolabeling, slides were incubated with DAPI (5min 1:5000 in PBS), washed (PBS, 3×5min) and mounted with Anti-Fade Fluorescence Mounting Medium. For α-Bungarotoxin staining, slides were tempered (15min RT) and processed according to instructions using α-Bungarotoxin CF-568 (biotium 0006).

#### Hematoxylin-eosin staining

Slides were tempered (15min RT), immersed in distilled water, dehydrated in increasing concentrations of ethanol (till 100%) and rehydrated to distilled water before staining. Slides were incubated 5min in Eosin, washed with water, incubated 2min in Hematoxylin, washed with water, dehydrated in increasing concentrations of ethanol (till 100%), incubated 3min in xylol, mounted in Eukitt (Biosystems) and dried 24h.

### Golgi-Cox staining

Dissected fresh brain was cut coronally in 3 parts and processed according to instructions (FD Rapid GolgiStain kit, PK401). PolyFreeze tissue freezing medium (Polysciences 19636-1) was used to cryopreserve pieces. Cerebellum was cut sagittally (300µm thickness; Leica cryostat) and processed according to instructions.

### Imaging and quantifications

Images of stained muscles were taken on an OLYMPUS microscope; cross-sectional areas and minimum Feret diameter were obtained with ImageJ. Fluorescence images were taken on a ZEISS Axioscan Z1 and a confocal LSM780 microscope (ZEN software). Acquisition parameters were optimized at beginning of experiment for each antibody (including for α-Bungarotoxin) and held constant. Maximal intensity z-projection of motor endplates (α-Bungarotoxin staining) was used for blind analysis of area, perimeter and intensity of staining using ImageJ. Maximal intensity z-projection was done with the same number of slides (same z-thickness) for brain. Thickness from granular and molecular layers of cerebellum was measured using ImageJ, along one comparable cerebellum slice per animal, and average value from each animal was used for the analysis between genotypes. Images of dendritic spines were taken from terminal dendrites of Purkinje neurons using an OLYMPUS microscope. Number of spines per length, spine head diameter and spine length were measured using ImageJ to evaluate spine density and spine morphology.

### Behavioral tests

#### Locomotion, rearing supported and unsupported in an Open Field (OF)

The same test and procedure described in ^12^ for grooming behavior were used for rearing supported (standing on hind limbs and one or two superior extremities) and unsupported (no superior limb used). Average velocity, total distance moved and time spent moving or not moving (considering thresholds of 2cm/s and 1.75cm/s, respectively) were calculated from the same recordings using EthoVision tracking software.

#### Circular Corridor (CC)

Rats were individually placed for 30min in a black acrylic circular corridor with external and internal diameters of 50cm and 40cm respectively, and a height of 40cm. Average velocity, total distance moved and time spent moving or not moving (considering thresholds of 2cm/s and 1.75cm/s, respectively) were calculated using EthoVision tracking software.

### Statistical analysis

Statistical analysis and graphs were conducted with R-3.5.1^28^. Shapiro test was used to assess normality of each sample. To address significance between WT and KI groups, Mann-Whitney tests were performed when normality was rejected, while for normal distributions, we used Bartlett tests to evaluate equality of variances and t-tests for equality of means. 2-way ANOVA was used for the analyses of gene expression and mTOR readouts (significant *P*-values from Tukey post-hoc analysis are reported in the figures). Generalized linear mixed models blocking litter as random factor were used for behavioral tests analysis (package lme4^29^), and obtained coefficients (mean and standard error of the mean) were used for graphs. Graphs were done using ggplot2 package^30^. *P*-values are reported in figure legends, statistical significance considered at *P*<0.05.

## RESULTS

### Plasmatic and urinary Crn is strongly reduced in *Slc6a8*^*xY389C/y*^ males

As previously shown^12^, *Slc6a8*^*xY389C/y*^ males exhibited the increased urinary Cr/Crn ratio marker of CTD. This increase was due to a significant increase in urinary Cr concentration via CrT impairment in kidney and to a strong significant reduction in urinary Crn levels (**Figure 1A** left). As Crn reuptake from primary urine is negligible, we asked whether plasmatic Crn levels were also decreased in KI males. As expected, plasmatic Crn levels in KI males were strongly reduced in comparison with those of WT males (**Figure 1A** middle), while plasmatic total CK only tended to decrease (**Figure 1A**, right).

**Figure 1:**
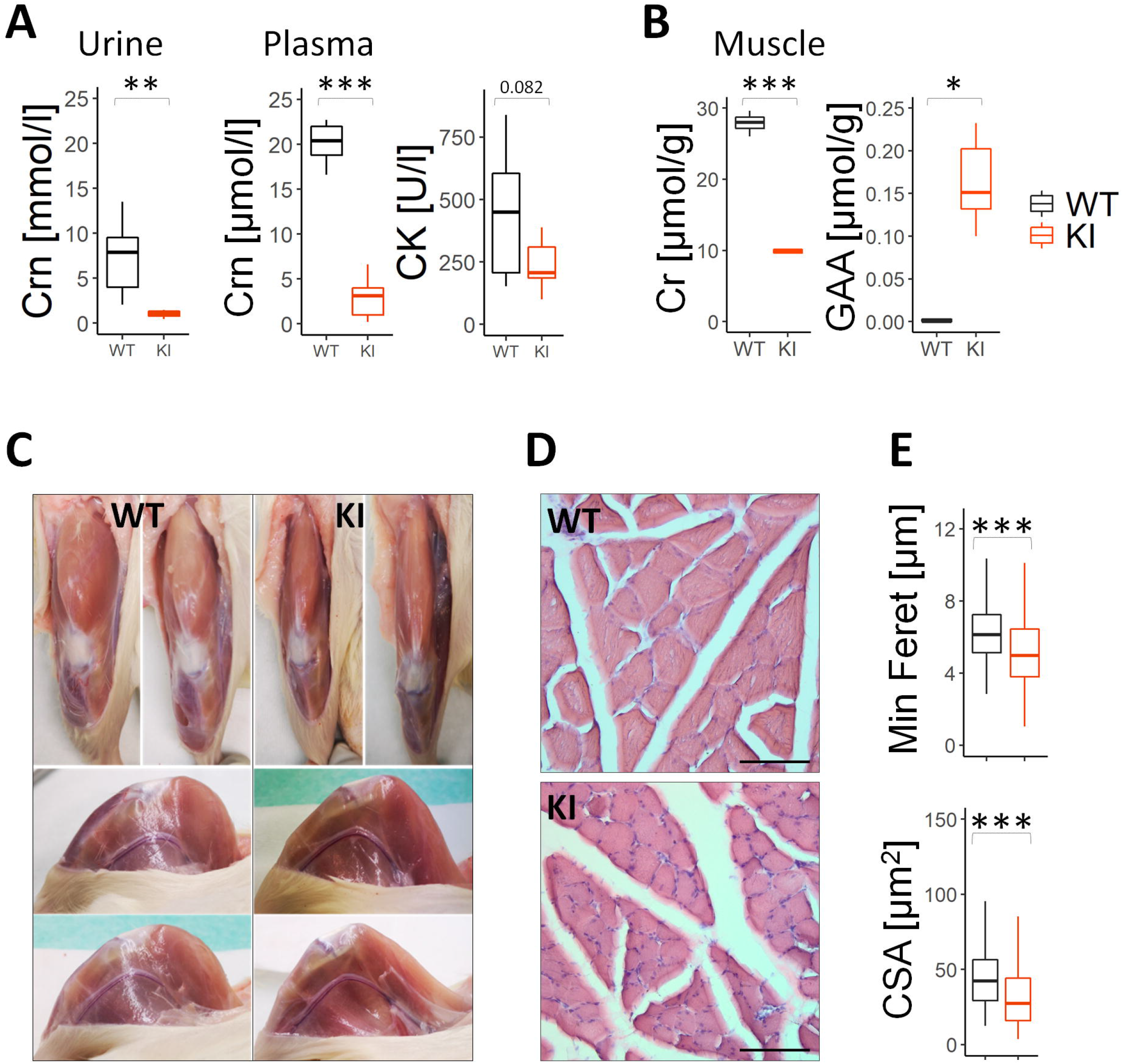
*S1c6a8*^*xY389C/y*^ KI males present reduced muscular mass and smaller myocytes. **A:** Urinary Crn (left panel) as well as plasmatic Crn and total CK levels (middle and right panel, respectively) as indicative of the muscular mass. 6WT and 6KI; two-tail t-test. **B:** Significant decrease of Cr ([µmol/g tissue], two-tail t-test) and significant increase of GAA levels in muscle ([µmol/g tissue], Mann-Whitney test) in KI male rats. 5WT and 5KI. **C:** Representative macroscopic pictures of WT and KI male hind limbs in frontal/longitudinal (upper panels) and medial/inner lateral views (lower panels) showing reduced volume of quadriceps in KI male rats. **D:** Representative microscopic pictures of hematoxylin/eosin staining in transversal sections of WT and KI male quadriceps showing significant smaller myocyte diameter in KI males. Scale bar=15µm. **E**: Quantifications of myocyte minimum Feret diameter and cross-sectional area per each genotype. Mann-Whitney test, 3WT and 3KI (319-382 measurements per WT and 414-555 per KI male). \**P*<0.05, \*\**P*<0.01, \*\*\**P*<0.001. Statistical analysis was conducted with R-3.5.1 ^28^. Graphs were done using ggplot2 package ^30^.

### Muscle from *Slc6a8*^*xY389C/y*^ males is Cr-deficient and increases its GAA content

Muscle is the main storage of peripheral Cr, and Crn is the spontaneous breakdown product of Cr. Therefore, low plasmatic and urinary Crn levels indicate low concentration of Cr in muscle and/or low muscular mass^31^. Cr levels were significantly reduced in *Slc6a8*^*xY389C/y*^ males muscles (**Figure 1B**, left). Interestingly, GAA levels in WT males muscles were almost undetectable but they were significantly increased in *Slc6a8*^*xY389C/y*^ males muscle (**Figure 1B**, right).

### *Slc6a8*^*xY389C/y*^ males exhibit reduced muscular mass and smaller myocytes

We explored quadriceps muscle to evaluate muscular mass. *Slc6a8*^*xY389C/y*^ males quadriceps were qualitatively thinner than those from WT (**Figure 1C**, note the rounded forms in WT *versus* KI males). In addition, myocyte minimum Feret diameter and myocyte cross-sectional area from *Slc6a8*^*xY389C/y*^ males were significantly reduced in comparison with those of WT males (**Figure 1D-E**). Neither vacuolization nor centralized nuclei were found in KI males muscles.

### Muscle from *Slc6a8*^*xY389C/y*^ males show no sign of muscle atrophy

Reduced muscular mass and myocyte size can indicate muscle atrophy^32^, which was reported in two CTD mouse models^11,33^ although no case was described in CTD patients so far. We therefore wondered whether *Slc6a8*^*xY389C/y*^ males present muscle atrophy. Muscle atrophy occurs when synthesis and degradation dynamic balance shifts towards protein degradation in response to several stimuli (e.g. infection, inflammation, oxidative or biomechanical stress)^32^. Such protein degradation is mainly mediated by the ubiquitin-proteasome system and autophagy via mTOR inhibition. FOXO-dependent genes, such as those encoding for E3 ubiquitin-ligases and autophagy-related proteins, are involved. We analyzed the expression of Atrogin-1 (an E3 considered a “master regulator” of muscle atrophy^34^), Itch/Aip4 (another E3 negatively regulating hypertrophy^35^), Gabarpl1/Gec1/Atg8l (implicated in autophagy^36^), and the mTOR-targeted proteins pS6K and pRPS6.

No significant differences between genotypes were found in the expression of atrogenes, ubiquitin-ligases or autophagy-related genes (**Figure 2A and Supplementary Figure 1**, using two different normalizations). In contrast, Redd1, a stress response gene also involved in mTOR inhibition^37^, was significantly increased in KI male muscles. Moreover, similar levels of the mTOR activation readouts, phosphorylated S6K1 (pS6K1) and RPS6 (pRPS6) over total S6K1 and RPS6, respectively, were found in muscles from both genotypes (**Figures 2B-C**). Intriguingly, levels of total and phosphorylated RPS6 (a substrate of S6K1 and other kinases^38^) but not those of S6K1 were significantly increased in KI male muscles (**Figures 2B-C**).

**Figure 2:**
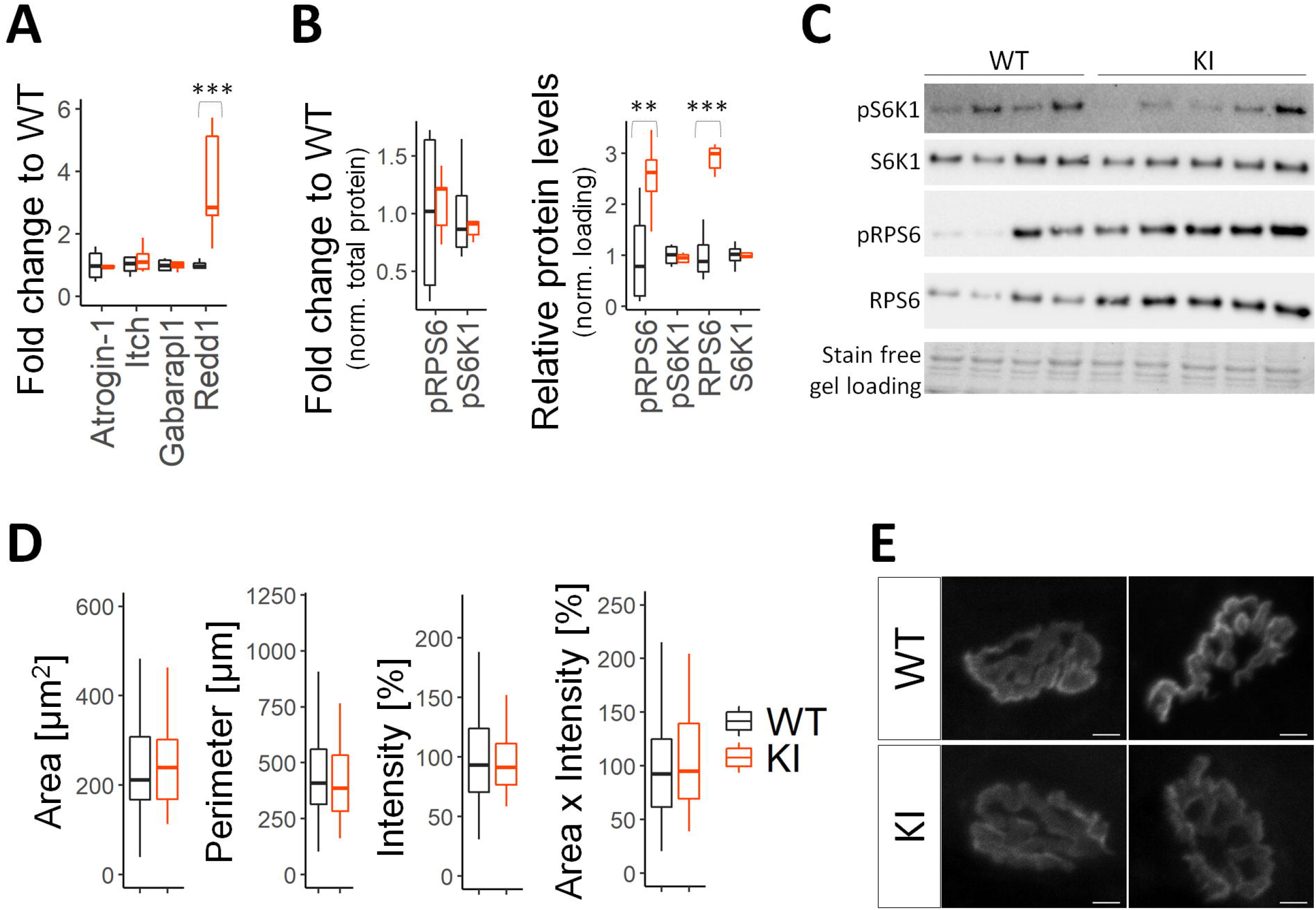
*Slc6a8*^*xY389C/y*^ KI males do not show consistent signs of muscle atrophy. **A:** Muscle expression levels (by quantitative PCR) of Atrogin-1, Itch and Gabarapl1 were similar between genotypes, and those of Redd1 significantly increased in muscle from KI males in comparison with those of WT males (orange and black boxes, respectively). 18S was used as housekeeping gene for normalization. 4 WT and 5 KI; 2-way ANOVA and Tukey posthoc test. **B:** Relative levels of phosphorylated S6K1 (pS6K1) and RPS6 (pRPS6) over total S6K1 and RPS6, respectively, were similar between genotypes (left panel). Relative total and phosphorylated protein levels of RPS6 were significantly increased in muscle from KI males while those of S6K1 were similar between genotypes (right panel). 4 WT and 5 KI; 2-way ANOVA and Tukey posthoc test. **C:** Representative western-blot from muscle of WT and KI males. **D:** Boxplots showing no significant differences between genotypes in the area (in µm^2^), perimeter (in µm), intensity (in % from WT mean value) or the combination of area and intensity (area multiplied by intensity, in % from WT mean value) of individual endplates stained with α-Bungarotoxin from muscle of WT and KI males. 3 WT and 3 KI using around 35 pictures per animal; Mann-Whitney test. **E:** Representative z-projection confocal images of individual endplates from WT and KI males using α-Bungarotoxin staining. Scale bar is 5 µm. \*\**P*<0.01, \*\*\**P*<0.001. Statistical analysis was conducted with R-3.5.1 ^28^. Graphs were done using ggplot2 package ^30^.

On the other hand, motor endplate (the specialized postsynaptic region of muscle cells) is reported to be structurally altered in cases of muscle atrophy such as sarcopenia^39^. We checked it by using the well-established α-Bungarotoxin staining taking advantage of its binding to nicotinic ACh receptors on motor endplates. No apparent endplate fragmentation was observed in muscles from both genotypes; area, perimeter and intensity of the staining were similar in muscles from both genotypes (**Figure 2D-E**).

These results show that *Slc6a8*^*xY389C/y*^ male muscles do not present signs of an atrophic process, although muscle cells appear to respond to some kind of stress.

### Motor function is affected in *Slc6a8*^*xY389C/y*^ males

Motor dysfunction is mentioned in 58% of CTD patients^40^. The most prevalent symptom is hypotonia, while other observations are signs of spasticity, coordination dysfunction (such as unstable gait or clumsiness) and dystonia. In order to assess motor function, we first checked signs of hypotonia by physical examination, then analyzed free exploration locomotion (using OF and CC tests) and rearing (spontaneous behavior requiring muscle performance, coordination and stability for its execution). Rearing (standing up with hind limbs) is “supported” if one or both superior limbs are used or “unsupported” when no superior limbs are used.

*Slc6a8*^*xY389C/y*^ males exhibited reduced body tension suggesting hypotonia, and frequent kyphosis or kyphoscoliosis worsening with aging (data not shown) suggesting muscle weakness. Additionally, *Slc6a8*^*xY389C/y*^ males showed a tendency to move less distance with less velocity in OF (**Figure 3A**) and moved significantly less distance with significantly less velocity, spending significantly less time moving in CC (**Figure 3A**). A more detailed analysis along time showed that these differences between genotypes started from minute 7 for cumulative distance moved and average velocity, and from minute 1 for cumulative time moving in CC test, while no difference along time was found in OF test (**Supplementary Figure 2**). Moreover, while *Slc6a8*^*xY389C/y*^ males showed a significant decrease in unsupported rearing (**Figure 3B**,), they did not show clear differences with WT males in supported rearing (**Figure 3C**). Furthermore, *Slc6a8*^*xY389C/y*^ males appeared to be clumsy, being less dynamic and more instable (more tensed, sometimes vibrating) in equilibrium postures and falling down sporadically. These results indicate altered motor function in *Slc6a8*^*xY389C/y*^ males.

**Figure 3:**
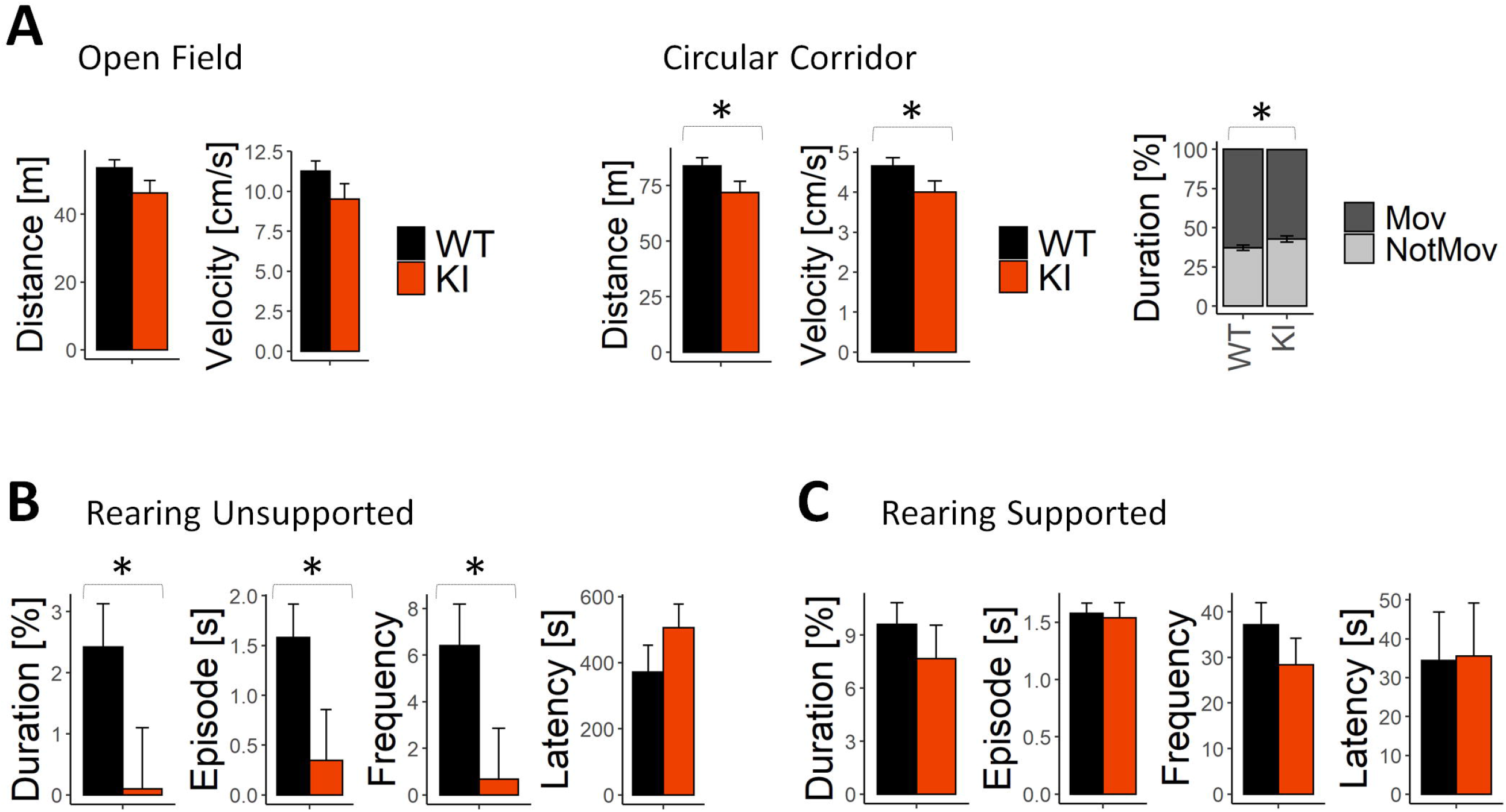
Motor function is affected in *Slc6a8*^*xY389C/y*^ KI males. **A:** In open field (OF) test, KI males (orange bars) tended to move less distance with less velocity in comparison with WT males (black bars); in circular corridor (CC) test, KI males moved significantly less distance with less velocity and spent significantly less time moving (Mov) than WT males (gray bars). 10 WT and 10 KI males. **B-C:** KI males showed significant differences in unsupported (no superior limbs used, **B**) but not in supported (**C**) rearing. 9 WT and 10 KI males. Linear mixed models blocking litter as random factor. \**P*<0.05. Statistical analysis was conducted with R-3.5.1 ^28^, package lme4 ^29^. Graphs were done using ggplot2 package ^30^.

### *Slc6a8*^*xY389C/y*^ male cerebellum is altered

Motor function depends on muscle and brain functions. One of CNS regions involved in motor function is cerebellum. Cerebellum presents very high levels of Cr in WT males and its Cr content is strongly decreased in *Slc6a8*^*xY389C/y*^ males^12^. We thus analyzed whether *Slc6a8*^*xY389C/y*^ male cerebellum was affected by Cr deficiency. Immunostainings of MAP2 (microtubule-associated protein 2, labeling neuronal dendrites and soma^41^) and NF-M/pNF-M (neurofilament mainly present in neuronal soma when dephosphorylated or in axons when phosphorylated^22,42^), showed decreased staining in both cerebellar granular and molecular layers from *Slc6a8*^*xY389C/y*^ males (**Figure 4A**). However, no difference was observed in NeuN immunostaining (**Figure 4B**), where the number of NeuN-positive cells (neurons) did not change significantly between genotypes (data not shown).

**Figure 4:**
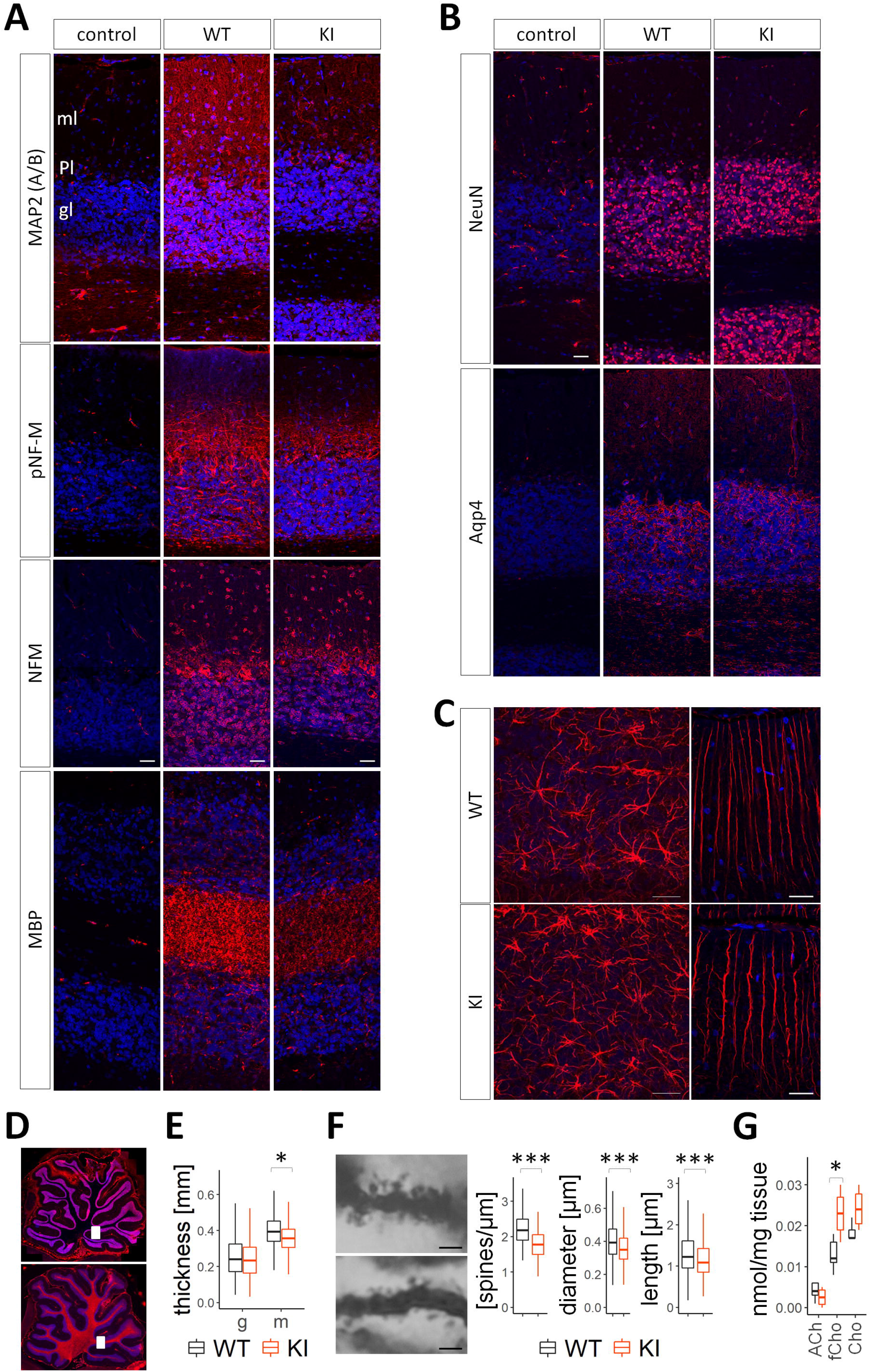
Cerebellum from *Slc6a8*^*xY389C/y*^ KI males is affected. **A:** Representative confocal z-projection images of MAP2 (A/B), pNF-M, NF-M and MBP (in red, DAPI in blue) showing decreased immunostaining in KI males. **B:** Representative confocal z-projection images of NeuN and Aqp4 immunostainings showing no differences between WT and KI males. **C:** Representative confocal images of GFAP immunostaining from granular layer (z-projection, left pictures) and molecular layer (slice of 0.30µm thickness, right pictures). **D:** Epifluorescence pictures from cerebellum in sagittal sections showing with white squares the regions in which confocal images were taken. All images correspond to the region labeled in the top picture except the one for MBP immunostaining (in **A**) which corresponds to the region labeled in the bottom picture. **E:** Quantification of molecular and granular layers thickness showing a significant reduction in the molecular (m) but not the granular (g) layer of KI males. Mann-Whitney test, \**P*<0.05, 4 WT and 4 KI. Average values from each animal were used for the analysis between genotypes (details in methods), plots showing the variability of all measurements. **F:** Representative images of Purkinje terminal dendrites from WT and KI males (top and bottom panels, respectively), and quantifications of spines density (left, 101 and 72 measurements from WT and KI respectively), head diameter (middle, 2024 and 1322 spines from WT and KI respectively) and spine length (right plot, 2368 and 1491 spines from WT and KI respectively) showing a significant reduction of the three measurements in KI males. Mann-Whitney test, 4 WT and 3 KI (3-4 neurons per animal; 20-30 measurements per rat for spine density, 410-620 spines per rat for head diameter and spine length), \*\*\**P*<0.001. **G:** Concentration of acetylcholine (Ach), free and total choline (fCho and Cho, respectively) in muscle from WT and KI males. 5 WT and 4 KI, t-test,\**P*<0.05. Calibration/scale bar in **A, B** and **C** represents 30µm; 2µm in **F**. Statistical analysis was conducted with R-3.5.1 ^28^. Graphs were done using ggplot2 package ^30^.

Myelin basic protein (MBP) immunostaining was patchy and decreased in *Slc6a8*^*xY389C/y*^ males (**Figure 4A**). Immunostaining of Aqp4, a protein highly expressed on perimicrovessel astrocytic foot processes (blood-brain barrier), glia limitans and ependyma^43^, was similar among cerebellums from both genotypes (**Figure 4B**). Nevertheless, GFAP immunostaining showed that astrocytes appeared more numerous in granular layer (**Figure 4C** left) while Bergmann glia processes were more tortuous in *Slc6a8*^*xY389C/y*^ males (**Figure 4C** right). Moreover, despite the variability in molecular and granular layers thickness along cerebellum, the molecular (but not the granular) layer was significantly thinner in *Slc6a8*^*xY389C/y*^ males (**Figure 4E**).

To figure out whether these changes might have an impact in neuronal function, we analyzed dendritic spines of terminal dendrites from Purkinje neurons. Purkinje neurons are some of the largest neurons in CNS, being the sole output from cerebellar cortex with important roles in motor coordination^44^. Spines are the postsynaptic compartment of excitatory synapses and their morphology correlates with synaptic strength. Changes in spine density and/or morphology could thus reflect synaptic alterations affecting neuronal function^45^. Purkinje neurons from *Slc6a8*^*xY389C/y*^ males showed significantly reduced spine density, and spines significantly thinner and shorter compared to WT (**Figure 4F**).

In summary, these results show that neurons, oligodendrocytes and astrocytes are affected in *Slc6a8*^*xY389C/y*^ males cerebellum, and together point towards altered morphology and connectivity which might have impacts in cerebellar function.

### Acetylcholine levels do not change in motor neurons from KI males

Other important nervous actors involved in motor function are motor neurons. They innervate muscle via cholinergic neurotransmission in a specialized synapse called neuromuscular junction. Given their role in several muscular disorders^46^, we wondered whether neurotransmission from motor neurons to muscle was affected in CTD. We evaluated ACh levels in muscle tissue, where motor neuron axon terminals are localized. In KI male muscles, ACh levels did not change significantly but tended to decrease in comparison with those of WT males (−37%, **Figure 4G**). Nevertheless, levels of its precursor, choline, were significantly increased (+77%, **Figure 4G**). These data reveal that final levels of ACh neurotransmitter tends to remain constant while choline homeostasis appears affected in KI males muscles.

## DISCUSSION

We describe the muscle and motor phenotype of the *Slc6a8*^*xY389C/y*^ CTD rat model. Brain Cr-deficient *Slc6a8*^*xY389C/y*^ rat males are also muscle Cr-deficient and present less muscle bulk without signs of atrophy, together with impaired motor function and morphological changes in cerebellum. Reduced muscle bulk of KI males is supported by reduced plasmatic and urinary Crn levels (muscular mass markers^47^) and thinner quadriceps. These results are in line with CTD patients, frequently showing reduced muscular mass^40^ and decreased plasmatic Crn levels^8^. Additionally, *Slc6a8*^*xY389C/y*^ male muscles presented decreased fiber size and Cr concentration (−64%). Similar muscle phenotype was described in two CrT KO mouse lines^11,33^ and was considered “muscle atrophy” despite alternative causes. One of these mouse lines showed strong histological deterioration of muscles in middle-age adults^33^, strongly suggesting atrophy. This was however not the case at earlier ages, and was not observed either in the other CrT KO mouse line^11^ nor in the KI rat males (even at 6 months-old, data not shown). We thus further explored whether the observed muscle phenotype was reflecting atrophy, and our results ruled it out: KI male muscles did not show increased expression of atrophy-involved genes, nor changes in mTOR signaling or in endplate morphology. This may also be true for the abovementioned CrT KO lines, at least at earlier ages. Such absence of atrophic process in KI male muscles is in agreement with the absence of reported muscular atrophy in CTD patients (no report of muscle atrophy, no muscle structural alterations in the sole analyzed for this^25^).

Reduced muscle bulk and smaller myocyte size in CTD rodent models are in agreement. This may be explained by the decreased muscle Cr concentration, observed in all ubiquitous CTD rodent models^9-12^, and the Cr roles in cell volume^2^ and muscle development^24^. Unaltered mTOR signaling suggests no disturbance in cell growth, but we cannot discard modifications in other pathways related with muscle growth and differentiation. Nevertheless, although reduced muscle mass is shared between CTD patients and rodent models, patients’ myocyte size has not been analyzed yet, and the unique CTD patient analyzed exhibited normal muscle Cr levels^25^. However, it is possible that other muscles from the same patient were Cr-deficient. Indeed, muscle Cr concentration can fluctuate depending on muscle type under Cr restriction, as observed in vegetarians^48^. Additional studies are needed to better characterize muscle phenotype in CTD patients, and to unravel the mechanisms leading to reduced muscular mass.

Noteworthy, two signals suggest that KI muscle cells are responding to some kind of stress: increased Redd1 as well as total and phosphorylated RPS6. Further investigations are needed to disentangle the mechanisms involved, which may represent potential therapeutic targets to prevent CTD muscle phenotype, or treat other muscle conditions, pathological or in the sports field.

We also analyzed plasmatic CK level as qualitative marker of skeletal muscle microtrauma^47^. *Slc6a8*^*xY389C/y*^ males showed low values but within normal range, in line with CTD patients^8^. This does not support muscle damage, being in agreement with absence of centrally nucleated myofibers, but suggests a reduction in muscle CK content, and by extension in Cr content as indeed observed in KI males.

Plasmatic and muscle Cr levels are both decreased in CTD rodents but both appear normal in patients^8,9,12^. Whether this difference is due to diet (vegetarian diet used in some CTD models^11,12^, while patients likely follow omnivorous diets) in combination or not with intrinsic factors (distinct regulation among species of Cr synthesis and transport, including alternative Cr transporters) is still unknown. Strikingly, this parallelism between muscle and plasmatic Cr levels is observed in both humans and rodents where synthesis and/or transport of Cr is affected^8,9,12,49,50^, as well as under Cr supplementation in conditions of Cr restriction with normal CrT function^48,50,51^. Plasmatic Cr might therefore be a marker of muscle Cr concentration, at least under possible conditions of Cr depletion.

Cr deficiency co-occurred with increased GAA concentration in *Slc6a8*^*xY389C/y*^ male muscles. Despite no available data from CTD patients, this is similar to another CrT KO mouse^14^. Such augmentation could indicate an attempt to compensate Cr deficiency by increasing *de novo* synthesis in muscle. Indeed, AGAT increased in muscles from CrT KO mice^11,52^ and in conditions of Cr depletion in kidney^53^.

Together with increased muscle GAA, KI males showed augmented plasmatic GAA^12^. Such pattern was also observed in GAMT deficiency^8,49^, suggesting that, similar to Cr, plasmatic GAA levels might inform about muscle GAA concentration. Nevertheless, in contrast with Cr being 3-times less concentrated in both plasma and muscle from KI *versus* WT males (**Table 2**), GAA in KI males increased by 126-times in muscle and only twice in plasma. This might reflect different tissular regulation of Cr synthesis and/or GAA transport (kidney-liver *versus* muscle). A conversion of muscle GAA concentration to µM (**Table 2**) allows the comparison between muscle and plasmatic GAA. Intriguingly, while WT males showed more GAA concentration in plasma than muscle, KI males presented the opposite. This dramatic GAA gradient change between plasma and muscle might have consequences in other peripheral tissues. However, whether muscle increases GAA uptake and/or synthesis or provides GAA to the rest of the body (thus being as important as kidney) in CTD conditions is not known.

**Table 2:** Comparison of metabolite concentrations [µmol/l] and their relative changes (%) in KI males with respect to those of WT males in urine, plasma and muscle. Average values (or median values if the distribution was not normal according to Shapiro test) per genotype of each metabolite concentration in urine, plasma and muscle. Concentration in muscle was transformed from µmol/g of tissue to **µmol/l** assuming that tissue contains 70% of water. Part of the data was published in ^12^. Significant differences between genotypes are recapitulated here and shown as \**P*<0.05, \*\**P*<0.01, \*\*\**P*<0.001.

|  | Urine |  | Plasma |  | Muscle |  | Relative change (%) |  |  |
| --- | --- | --- | --- | --- | --- | --- | --- | --- | --- |
|  | WT | KI | WT | KI | WT | KI | urine | plasma | muscle |
| **Crn** | 7340 | 950 | 20.15 | 2.65 | NA | NA | 12.9 ** | 13.2 *** | NA |
| **GAA** | 620 | 1300 | 5 | 10 | 0.84 | 105.7 | 210.3 | 196.5 ** | 12583 * |
| **Cr** | 280 | 9535 | 220 | 63 | 28000 | 10000 | 12109 ** | 32.2 *** | 36.0 *** |

Cr is essential for muscle strength and performance^54^. Low muscle Cr content coupled to reduced muscular mass and myocyte size points towards low muscle performance likely affecting motor function. Indeed, *Slc6a8*^*xY389C/y*^ males exhibited signs of lower muscle tone (hypotonia) and muscle weakness in early adulthood, and spent less time moving in comparison with WT males (CC test). Similar phenotype was found in a CTD mouse model^11^ and, although muscle weakness appears only occasionally in CTD patients, hypotonia is frequent^8^. Moreover, KI males showed other signs of motor dysfunction such as reduced locomotion and unsupported rearing. The differences in distance moved and velocity only appeared with time in CC test, suggesting a relation with reduced muscle performance although we cannot discard dysfunctions from CNS. On the other hand, *Slc6a8*^*xY389C/y*^ males accomplished supported but not unsupported rearing as WT males, revealing a not too severe or disabling phenotype with extra difficulties involving muscle strength, stability and/or coordination. These motor dysfunction symptoms are in line with those of CTD patients^8,40^ and a mouse model^11^. They can be due to altered muscle and/or CNS functions, although each one’s contribution to this phenotype is not easy to detangle.

CNS alterations have been described in CTD patients and mouse models^8,9,14,16^. However, cerebellum, a brain region with high Cr content involved in coordination and balance and with abnormalities in CTD patients, has been very little explored so far. The Cr-deficient KI males cerebellum exhibited similar number of NeuN-positive cells but changes in neuronal cytoskeleton-related proteins, suggesting that the number of neurons did not change although both dendritic trees and axons might be altered in *Slc6a8*^*xY389C/y*^ males cerebellum. This, together with decreased MBP staining, in agreement with delayed myelination of some patients^8^, suggests that neuronal connectivity might be affected under CTD^21,22^.

Interestingly, differences in GFAP staining indicate disturbed astrocytes in *Slc6a8*^*xY389C/y*^ males despite not being direct CTD targets since they do not usually express CrT^55^. Bergmann glia, specialized astrocytes playing roles in early cerebellar development (granule cells migration^56^), required for synaptic pruning and intimately associated with Purkinje cells^57,58^, had tortuous processes in *Slc6a8*^*xY389C/y*^ males. This, together with the observed changes in molecular layer (reduced thickness and MAP2 staining), may reflect altered Purkinje cells (the sole output from cerebellar cortex), particularly in their dendritic tree. To see whether these alterations may impact neuronal function, we analyzed Purkinje neurons’ dendritic spines. Morphology of these excitatory postsynaptic compartments correlates with synaptic strength^45^. Strikingly, Purkinje neurons from KI males showed decreased spine density and size, indicating that Purkinje neurons might present less excitatory inputs and lower synaptic strength resulting in decreased signaling^59^, which could alter cerebellar function and in turn lead to motor dysfunction. These limitations in number and size of dendritic spines could be due to either neuronal network modifications triggering changes in connectivity and/or to intrinsic structural and metabolic alterations linked to Cr deficiency, likely including a compromised cell energy state.

We show for the first time alterations in cerebellum from a CTD animal model. We demonstrated that Cr-deficient *Slc6a8*^*xY389C/y*^ male cerebellum^12^ present morphological changes which are in line with abnormalities seen in CTD patients^8^. Noteworthy, we demonstrate that *in vivo* Cr deficiency affects cerebellar neurons, astrocytes and oligodendrocytes and alters Purkinje cell spines, limiting their number and size. These morphological changes could impact cerebellar function contributing to the *Slc6a8*^*xY389C/y*^ males motor dysfunction phenotype.

We also explored whether motor neurons were involved in this motor function impairment. ACh, only attributable to motor neuron terminals in muscle tissue, remained constant (although tending to decrease) in KI males. This result, together with the similar staining of ACh receptors at endplates, suggests that cholinergic neurotransmission is not affected in KI males muscle, at least in young animals.

On the other hand, free choline in muscle is present in motor neurons and other cells, such as myocytes. Choline is also precursor for choline phospholipids and S-adenosyl-methionine, important for methylation of molecules such as GAA for Cr synthesis. Such increase in choline concentration co-occurred with increased GAA concentration, suggesting that these changes might be related with the possible attempt in increasing Cr levels in KI males muscle.

In summary (**Table 3**), we characterized the motor function phenotype of *Slc6a8*^*xY389C/y*^ rat males and provided more cues about CTD pathology and *in vivo* effects of Cr deficiency in muscle and cerebellum.

**Table 3:**
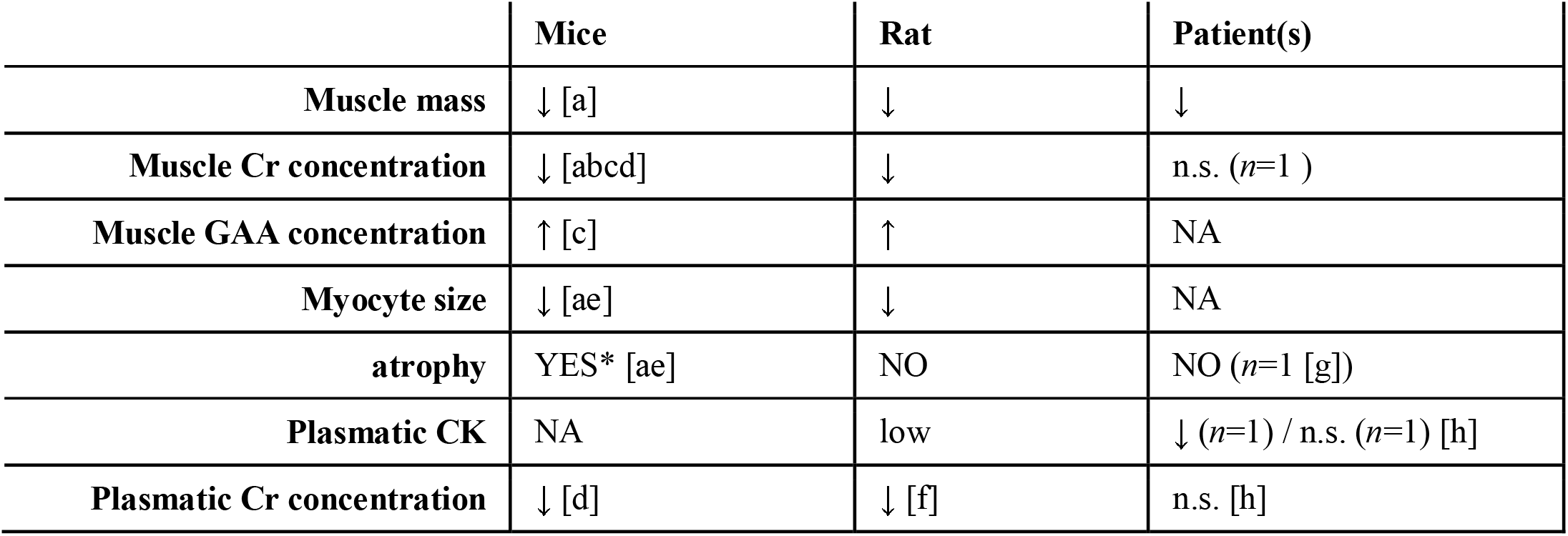
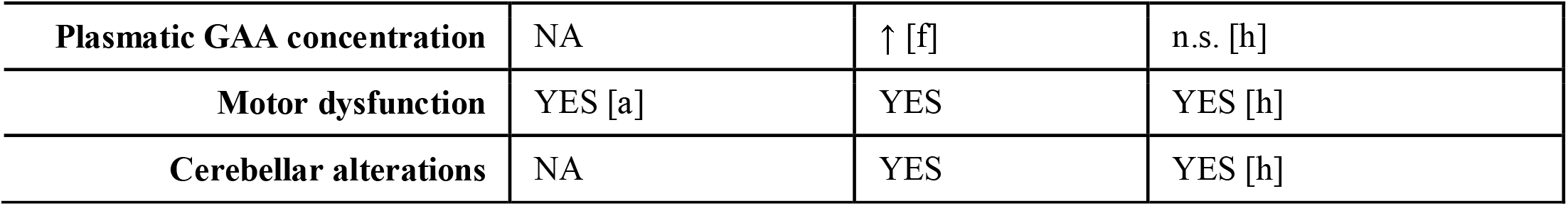
Comparisons of different features among rat and mouse models of CTD and CTD patients. ↓: significant decrease; ↑: significant increase; low: decreased but in normal range; n.s.: no significant differences; NA: not analyzed; YES: feature present; YES*: feature clearly present in one only at certain age ^33^ and mentioned but not fully demonstrated in other ^11^; NO: feature absent; a (Stockebrand et al., 2018); b, (Baroncelli et al., 2014); c, (Baroncelli et al., 2016); d, (Skelton et al., 2011); e (Wawro et al., 2021); f (Duran-Trio et al., 2021); g (Pyne-Geithman et al., 2004); h (van de Kamp et al., 2014).

|  | Mice | Rat | Patient(s) |
| --- | --- | --- | --- |
| **Muscle mass** | ↓ [a] | ↓ | ↓ |
| **Muscle Cr concentration** | ↓ [abcd] | ↓ | n.s. (n=1 ) |
| **Muscle GAA concentration** | ↑ [c] | ↑ | NA |
| **Myocyte size** | ↓ [ae] | ↓ | NA |
| **atrophy** | YES* [ae] | NO | NO (n=1 [g]) |
| **Plasmatic CK** | NA | low | ↓ (n=1) / n.s. (n=1) [h] |
| **Plasmatic Cr concentration** | ↓ [d] | ↓ [f] | n.s. [h] |
| **Plasmatic GAA concentration** | NA | ↑ [f] | n.s. [h] |
| **Motor dysfunction** | YES [a] | YES | YES [h] |
| **Cerebellar alterations** | NA | YES | YES [h] |

## ACKNOWLEDGEMENTS

This work was supported by the Swiss National Science Foundation (Grant n° 31003A-175778). We thank Marc Loup, Marc Lanzillo, Dario Sessa, Ana Versace and Annie-Juliette Ferrari-Gyger for excellent technical help. The authors declare no competing financial interests.

## AUTHOR CONTRIBUTIONS

LDT and OB designed the study and wrote the manuscript; LDT and GFP handled the animals and collected samples and liquids; CR and PAB performed Cr, GAA, Crn and CK measures; LDT performed acetylcholine and choline measures; LDT, JG and CS designed and performed the behavioral tests; experiments and analysis of gene expression and western blots were designed and performed by ISA and KDB; fluorescence and histological studies were performed by LDT; Golgi-Cox staining was performed by GFP and LDT and dendritic spine studies by LDT; LDT analyzed the data; all resources needed to perform this work were provided by OB, CC, CS and KDB; OB and CC secured the financial support of this work; all authors reviewed and corrected the manuscript.

