## Supplementary material for "Creatine transporter deficient rat model show motor dysfunction, cerebellar alterations and muscle creatine deficiency without muscle atrophy"

**METHODS**

**Detailed list of material and reagents used:**

| **Procedures** | **Reagents** |
| --- | --- |
| RT-qPCR | TRIzol (Thermo Fisher, 15596018) |
|  | mRNA extraction column (Thermo Fisher, 12183018A) |
|  | High Capacity cDNA Reverse Transcription Kit (Thermo Fisher, 4368813) |
|  | SYBER Green-based master mix (ThermoFisher Scientific, A25778) |
| WB | PVDF membrane (Bio rad 456 8086) |
|  | anti-phospho-S6K1 Thr389 (pS6K1, Cell signaling, 9234) |
|  | anti-phospho-S6 Ribosomal Protein Ser235/236 (pRPS6, Cell signaling, 2211) |
|  | anti-S6K1 (Cell signaling, 9202) |
|  | anti-S6 Ribosomal Protein (RPS6, Cell signaling, 2217) |
|  | anti-rabbit IgG HRP-linked antibodies (Cell signaling, 7074) |
| Histology and immunofluorescence | Tissue-Tek (OCT4583) |
|  | gelatin (Sigma-Aldrich G2500) |
|  | anti-MAP2(A/B), MAB378 |
|  | anti-pNF-M(NN18), MAB5254 |
|  | anti-NF-M (RM044), MAB5254 |
|  | anti-MBP(F6), sc-271524 |
|  | anti-NeuN, MAB377 |
|  | anti-GFAP, Abcam 7779-500 |
|  | anti-Aqp4 (C-term), AB3594 |
|  | goat anti-rabbit-A555, A21429 |
|  | goat anti-mouse-A555, A21422 |
|  | DAPI (Thermo Fisher D1306) |
|  | Anti-Fade Fluorescence Mounting Medium (ab104135) |
|  | α-Bungarotoxin CF-568 (biotium 0006) |
|  | Ehrlich Eosin (Fluka) |
|  | Harris Hematoxylin (Histolab) |
|  | Eukitt (Biosystems) |
| Golgi-Cox staining | FD Rapid GolgiStain kit, PK401 |
|  | PolyFreeze tissue freezing medium (Polysciences 19636-1) |

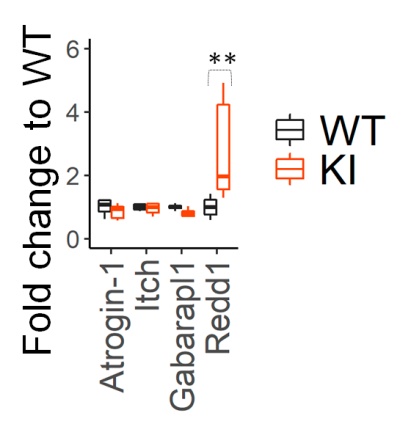

### RESULTS

**Supplementary Figure 1:** Muscle expression levels (by quantitative PCR) of Atrogin-1, Itch, Gabarapl1 and Redd1 from WT and KI males (black and orange boxes, respectively). GAPDH was used as housekeeping gene for normalization.

4 WT and 5 KI; 2-way ANOVA and Tukey posthoc test. ***P*<0.01, ****P*<0.001. Statistical analysis was conducted with R-3.5.1 ^28^. Graphs were done using ggplot2 package ^30^.

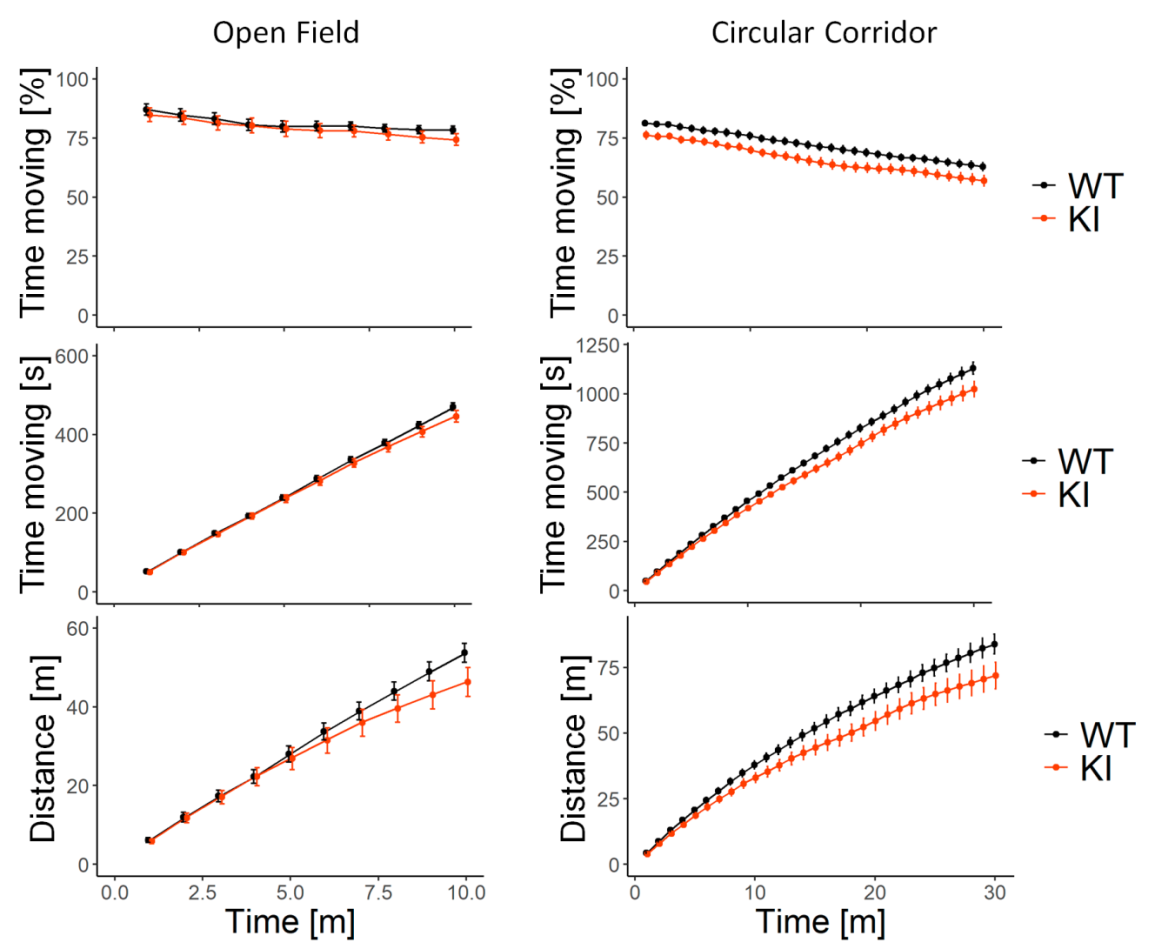

**Supplementary Figure 2: Cumulative time moving and cumulative distance along time in open field and circular corridor tests.**

No differences were found between genotypes in the open field test in cumulative time moving (in % and in seconds, upper and middle left panels respectively) or cumulative distance moved (in meters, bottom left panel) at any time. 10 WT and 10 KI males.

In circular corridor test, KI males spent significantly (*P*<0.05) less time moving from the first minute (upper and middle right panels) and moved significantly less distance from minute 7 (bottom right panel) in comparison with WT males. 9 WT and 10 KI males.

Linear mixed models blocking litter as random factor. Statistical analysis was conducted with R-3.5.1 ^28^, package lme4 ^29^. Graphs were done using ggplot2 package ^30^.
